# Gene model for the ortholog of *cnk* in *Drosophila erecta*

**DOI:** 10.1101/2025.08.06.668985

**Authors:** Leon F. Laskowski, Brianna Cowan, Greta Leissa, Harper O. W. Wallace, Indrani Bose, Jeffrey S. Thompson, Maria S. Santisteban

## Abstract

Gene model for the ortholog of *connector enhancer of ksr* (*cnk*) in the Dere_CAF1 Genome Assembly (GenBank Accession: GCA_000005135.1) of *Drosophila erecta*. This ortholog was characterized as part of a developing dataset to study the evolution of the Insulin/insulin-like growth factor signaling pathway (IIS) across the genus *Drosophila* using the Genomics Education Partnership gene annotation protocol for Course-based Undergraduate Research Experiences.

## Introduction

*This article reports a predicted gene model generated by undergraduate work using a structured gene model annotation protocol defined by the Genomics Education Partnership (GEP; thegep.org) for Course-based Undergraduate Research Experience (CURE). The following information in quotes may be repeated in other articles submitted by participants using the same GEP CURE protocol for annotating Drosophila species orthologs of Drosophila melanogaster genes in the insulin signaling pathway*.

“Computational gene predictions in non-model organisms often can be improved by careful manual annotation and curation, allowing for more accurate analyses of gene and genome evolution (Mudge and Harrow 2016; Tello-Ruiz et al., 2019) The Genomics Education Partnership (thegep.org) uses web-based tools to allow undergraduates to participate in course-based research by generating manual annotations of genes in non-model species (Rele et al., 2023). These models of orthologous genes across species, such as the one presented here, then provide a reliable basis for further evolutionary genomic analyses when made available to the scientific community.” (Myers et al., 2024).

“The particular gene ortholog described here *connector enhancer of ksr* (*cnk*) in *D. erecta* was characterized as part of a developing dataset to study the evolution of the Insulin/insulin-like growth factor signaling pathway (IIS) across the genus *Drosophila*. The insulin signaling pathway is a highly conserved pathway in animals and is central to nutrient uptake (Grewal 2009; Hietakangas and Cohen 2009).” (Myers et al., 2024).

“*Connector enhancer of ksr* (*cnk*, also known as CG6556; FBgn0286070) was identified in *Drosophila* by a screen for mutations that modify a *ksr* (kinase suppressor of RAS)-dependent phenotype (Therrien et al., 1998). *cnk* has diverse functions downstream of RTK (receptor tyrosine kinase) signaling events, including EGFR (epidermal growth factor)-mediated patterning of wing disc territories, and air sac development in the dorsal thorax (Baonza et al., 2000; Cabernard and Affolter 2005). The large multidomain Cnk protein functions as a molecular scaffold to recruit and integrate signaling components (Laberge et al., 2005; Clapéron and Therrien 2007; Wolfstetter et al., 2017). In addition to its scaffold function, Cnk appears to directly induce RAF (of the Ras-Raf-MAPK signaling pathway) catalytic function by a kinase-independent mechanism (Clapéron and Therrien 2007).” (Lawson et al., 2024).

“*D. erecta* is part of the *melanogaster* species group within the subgenus *Sophophora* of the genus *Drosophila* (Sturtevant 1939; Bock 1972). It was first described by Tsacas and Lachaise (1974). *D. erecta* is found in west central Africa (https://www.taxodros.uzh.ch/, accessed 17 Jully 2025; Markow and O’Grady 2006) where it is found to breed primarily on the fruits of *Pandanus candelabrum*, a spiny, evergreen shrub (Unwin 1920; Lachaise and Tsacas 1983).” (Lieser et al., 2024).

We propose a gene model for the *D. erecta* ortholog of the *D. melanogaster connector enhancer of ksr* (*cnk*) gene. The genomic region of the ortholog corresponds to the uncharacterized protein LOC6548136 (RefSeq accession XP_001975228.1) in the Dere_CAF1 Genome Assembly of *D. erecta* (GenBank Accession: GCA_000005135.1 - Drosophila 12 Genomes Consortium et al., 2007; Chirn et al., 2015; Ma et al., 2018). This model is based on RNA-Seq data from *D. erecta* (PRJNA414017, PRJNA264407) and *cnk* in *D. melanogaster* using FlyBase release FB2022_04 (GCA_000001215.4; Larkin et al., 2021; Gramates et al., 2022; Jenkins et al., 2022)

### Synteny

The target gene, *cnk*, occurs on chromosome 2R in *D. melanogaster* and is flanked upstream by *CG6550* and lethal (2) k01209 (*l(2)k01209*) and downstream by Proteasome α5 subunit (*Prosalpha5*) and *Vajk4*. The *tblastn* search of *D. melanogaster* cnk-PB (query) against the *D. erecta* (GenBank Accession: GCA_000005135.1) Genome Assembly (database) placed the putative ortholog of *cnk* within scaffold scaffold_4845 (CH954179.1) at locus LOC6548136 (XP_001975228.1)— with an E-value of 0.0 and a percent identity of 81.74%. Furthermore, the putative ortholog is flanked upstream by LOC6548134 (XP_001975226.1) and LOC6548135 (XP_001975227.1), which correspond to *CG6550* and *l(2)k01209* in *D. melanogaster* (E-value: 0.0 and 0.0; identity: 96.02% and 97.12%, respectively, as determined by *blastp*; Figure 1A, Altschul et al., 1990). The putative ortholog of *cnk* is flanked downstream by LOC6548137 (XP_001975229.3) and LOC6548138 (XP_001975230.1), which correspond to *Prosalpha5* and *Vajk4* in *D. melanogaster* (E-value: 9e-173 and 0.0; identity: 93.03% and 99.39%, respectively, as determined by *blastp*). The putative ortholog assignment for *cnk* in *D. erecta* is supported by the following evidence: The genes surrounding the *cnk* ortholog are orthologous to the genes at the same locus in *D. melanogaster* and synteny is completely conserved, supported by results generated from *blastp* (E-values and percent identities); we conclude that LOC6548136 is the correct ortholog of *cnk* in *D. erecta* (Figure 1A).

**Figure 1:**
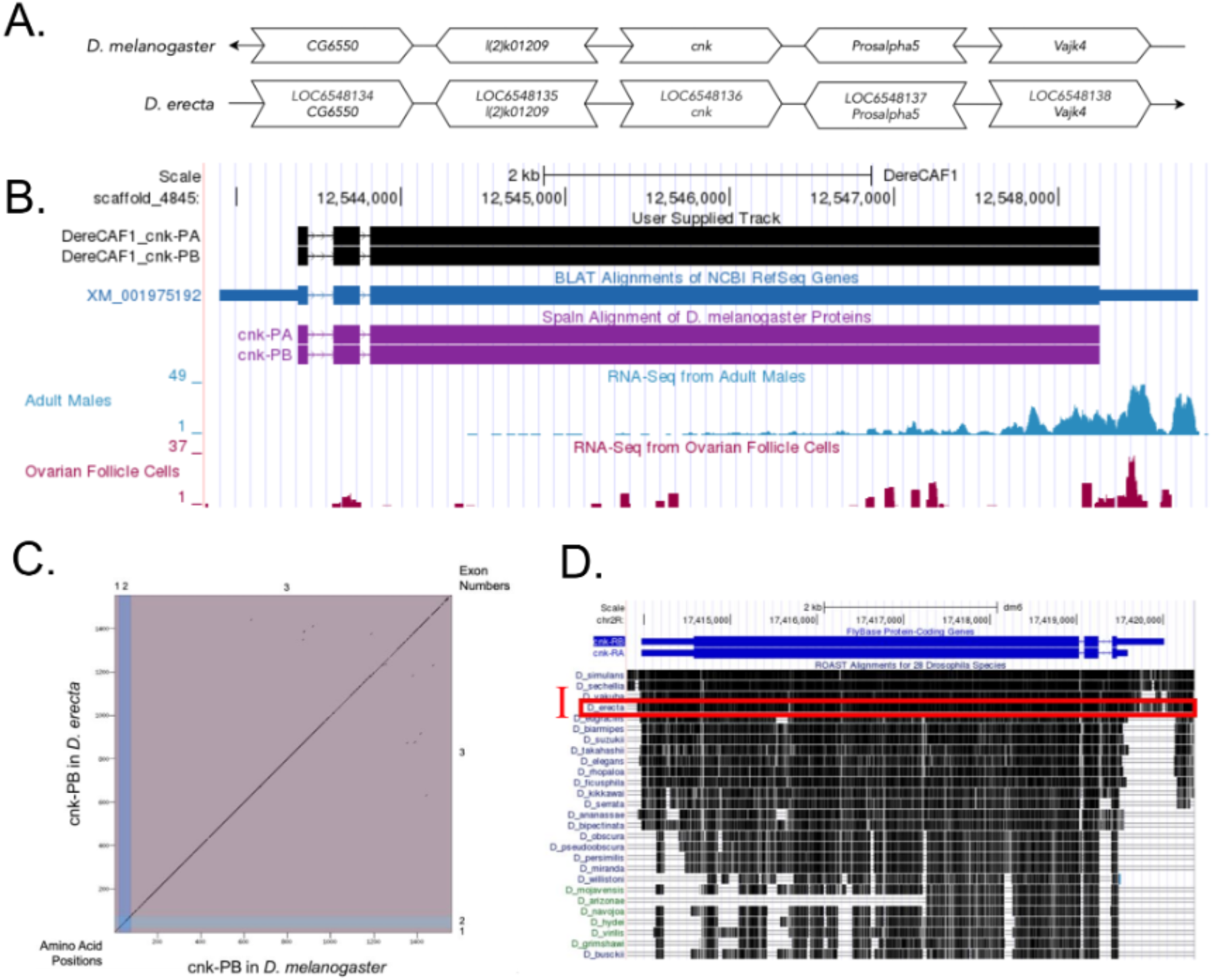
*cnk* gene model comparison between *Drosophila erecta* and *Drosophila melanogaster* orthologs. **(A) Synteny comparison of the genomic neighborhoods for *cnk* in *Drosophila melanogaster* and *D. erecta***. Thin underlying arrows indicate the DNA strand within which the target gene–*cnk*–is located in *D. melanogaster* (top) and *D. erecta* (bottom). The thin arrow pointing to the right indicates that *cnk* is on the positive (+) strand in *D. erecta*, and the thin arrow pointing to the left indicates that *cnk* is on the negative (-) strand in *D. melanogaster*. The wide gene arrows pointing in the same direction as *cnk* are on the same strand relative to the thin underlying arrows, while wide gene arrows pointing in the opposite direction of *cnk* are on the opposite strand relative to the thin underlying arrows. White gene arrows in *D. erecta* indicate orthology to the corresponding gene in *D. melanogaster*. Gene symbols given in the *D. erecta* gene arrows indicate the orthologous gene in *D. melanogaster*, while the locus identifiers are specific to *D. erecta*. **(B) Gene Model in GEP UCSC Track Data Hub (Raney et al**., **2014)**. The coding-regions of *cnk* in *D. erecta* are displayed in the User Supplied Track (black); CDSs are depicted by thick rectangles and introns by thin lines with arrows indicating the direction of transcription. Subsequent evidence tracks include BLAT Alignments of NCBI RefSeq Genes (dark blue, alignment of Ref-Seq genes for *D. erecta*), Spaln of D. melanogaster Proteins (purple, alignment of Ref-Seq proteins from *D. melanogaster*), RNA-Seq from Adult Males and Ovarian Follicle Cells (light blue and red respectively; alignment of Illumina RNA-Seq reads from *D. erecta*). **(C) Dot Plot of cnk-PB in *D. melanogaster* (*x*-axis) vs. the orthologous peptide in *D. erecta* (*y*-axis)**. Amino acid number is indicated along the left and bottom; CDS number is indicated along the top and right, and CDSs are also highlighted with alternating colors. **(D) Conservation of 28 *Drosophila* species showing evolutionary conservation of cnk-PB as shown in *D. melanogaster***. The black regions in the ROAST Alignments for 28 *Drosophila* Species indicate conservation relative to *D. melanogaster*. This region shows high local sequence similarity to *D. melanogaster* in multiple species. The ROAST alignment for *D. erecta* is highlighted by the red box denoted I.

### Protein Model

*cnk* in *D. erecta* has two identical protein-coding isoforms (cnk-PA and cnk-PB; Figure 1B). The isoforms contain three CDSs each. Relative to the ortholog in *D. melanogaster*, the CDS number and isoform count are conserved. The sequence of cnk-PB in *D. erecta* has 96.54% identity (E-value: 0.0) with the protein-coding isoform cnk-PB in *D. melanogaster*, as determined by *blastp* (Figure 1C). Differences were found in the level and type of RNA-seq data supporting this model. This gene model can be viewed in the *D. erecta* genome at this TrackHub.

### Special characteristics of the protein model

#### Insufficient RNA-seq data in *D. erecta* to support the *cnk* gene model

The May 2011 (Agencourt Dere_CAF1/DereCAF1) assembly in *D. erecta* has Adult Male and Ovarian Follicle Cell RNA-seq data. The current RNA-seq data found in *D. erecta* CAF1 assembly (PRJNA414017, PRJNA264407) shows discontinuous RNA-seq data within the coding sequence of *cnk*. However, the *Drosophila* Conservation of 28 species track in *D. melanogaster* indicates high identity in the coding sequence between *D. melanogaster* and *D. erecta* (Red box I, Figure 1D). However, long-read RNA sequence data is required to confirm the current gene model of *cnk* in *D. erecta*.

## Methods

Detailed methods including algorithms, database versions, and citations for the complete annotation process can be found in (Rele et al., 2023). Briefly, students use the GEP instance of the UCSC Genome Browser v.435 (https://gander.wustl.edu; Kent et al., 2002; Navarro Gonzalez et al., 2021) to examine the genomic neighborhood of their reference IIS gene in the *D. melanogaster* genome assembly (Aug. 2014; BDGP Release 6 + ISO1 MT/dm6). Students then retrieve the protein sequence for the *D. melanogaster* reference gene for a given isoform and run it using *tblastn* against their target *Drosophila* species genome assembly on the NCBI BLAST server (https://blast.ncbi.nlm.nih.gov/Blast.cgi; Altschul et al., 1990) to identify potential orthologs. To validate the potential ortholog, students compare the local genomic neighborhood of their potential ortholog with the genomic neighborhood of their reference gene in *D. melanogaster*. This local synteny analysis includes at minimum the two upstream and downstream genes relative to their putative ortholog. They also explore other sets of genomic evidence using multiple alignment tracks in the Genome Browser, including BLAT alignments of RefSeq Genes, Spaln alignment of *D. melanogaster* proteins, multiple gene prediction tracks (e.g., GeMoMa, Geneid, Augustus), and modENCODE RNA-Seq from the target species. Detailed explanation of how these lines of genomic evidenced are leveraged by students in gene model development are described in (Rele et al., 2023). Genomic structure information (e.g., CDSs, intron-exon number and boundaries, number of isoforms) for the *D. melanogaster* reference gene is retrieved through the Gene Record Finder (https://gander.wustl.edu/~wilson/dmelgenerecord/index.html; (Rele et al., 2023). Approximate splice sites within the target gene are determined using *tblastn* using the CDSs from the *D. melanogaste*r reference gene. Coordinates of CDSs are then refined by examining aligned modENCODE RNA-Seq data, and by applying paradigms of molecular biology such as identifying canonical splice site sequences and ensuring the maintenance of an open reading frame across hypothesized splice sites. Students then confirm the biological validity of their target gene model using the Gene Model Checker (https://gander.wustl.edu/~wilson/dmelgenerecord/index.html; (Rele et al., 2023), which compares the structure and translated sequence from their hypothesized target gene model against the *D. melanogaster* reference gene model. At least two independent models for a gene are generated by students under mentorship of their faculty course instructors. Those models are then reconciled by a third independent researcher mentored by the project leaders to produce the final model. Note: comparison of 5’ and 3’ UTR sequence information is not included in this GEP CURE protocol.

## Supporting information

Gene Model Data Files

## Supplemental Files

1. Zip file containing a FASTA, PEP, GFF files for the gene model
2. Figure 1 in high resolution

## Metadata

Bioinformatics, Genomics, *Drosophila*, Genotype Data, New Finding

## Acknowledgements

We would like to thank Wilson Leung for developing and maintaining the technological infrastructure that was used to create this gene model, Bethany C. Lieser for retrofitting this model and Laura K. Reed for overseeing the project. Thank you to FlyBase for providing the definitive database for *Drosophila melanogaster* gene models. FlyBase is supported by grants: NHGRI U41HG000739 and U24HG010859, UK Medical Research Council MR/W024233/1, NSF 2035515 and 2039324, BBSRC BB/T014008/1, and Wellcome Trust PLM13398.

